# Chromophore charge-state switching through copper-dependent homodimerisation of an engineered green fluorescent protein

**DOI:** 10.1101/2025.08.27.672602

**Authors:** Rochelle D Ahmed, Danoo Vitsupakorn, Kieran D. Hartwell, Karma Albalawi, Pierre Rizkallah, Peter D. Watson, D. Dafydd Jones

**Affiliations:** School of Biosciences, Molecular Biosciences Division, Cardiff University, Sir Martin Evans Building, Cardiff, CF10 3AX, UK; School of Medicine, Cardiff University, Cardiff, CF14 4XN, UK; Department of Chemistry, Faculty of Science, University of Tabuk, Tabuk, Saudi Arabia

## Abstract

Here, we have linked one of the most common protein-protein interaction events, homodimerisation, to an essential trace metal, copper, through engineering green fluorescent protein. Mutation of H148 to cysteine promotes the neutral chromophore in the monomer that excites predominantly at ∼400 nm. Homodimerisation via a copper-dependent disulphide bridge, switches the chromophore to the charged phenolate that excites at ∼490 nm. The result is ∼30 fold change in the fluorescence emission ratio. Homo-dimerisation kinetics are further improved by optimising the sfGFP homodimer interface, generating the variant termed GFP-diS2. Structures of the monomeric and dimeric GFP-diS2 suggests charge switching is through peptide bond flipping and the formation of a buried organised water networks around the chromophore that span the interface region. Fusion to a leucine zipper protein dimerisation element greatly increased GFP-diS2 association rate making it a more effective copper sensor in vitro and in vivo with Cu(I) instigating the signal change quicker and at lower ion concentrations than Cu(II). Thus, GFP-diS2 provides the framework for generating a sensitive genetically encoded copper sensor and will eventually be adapted to monitor one of the most important protein-protein interactions in biology, homo-oligomerisation.

## Introduction

Fluorescent proteins (FPs) have revolutionised how researchers investigate biological processes ^1–3^. Originally used as passive tags simply to fuse to a protein of interest, there has been a shift to monitoring active and dynamic processes that report on multi-component events ^3–5^. Protein-metal and protein-protein interactions (PPIs) are two such areas. Metals play a critical role in biology with an estimated 30-50% of proteins dependent on metals for their structure and/or function ^6, 7^. Copper is a particularly important trace metal in biology having roles ranging from enzyme catalysis, to signalling, to redox processes ^8–11^. Due to its role in the formation of reactive oxygen species, free copper ion concentration is tightly regulated in biological systems ^11^. Thus, the development of genetically encoded copper-responsive fluorescent proteins has been of significant interest ^5, 12, 13^. However, many current copper sensors have small responses, lack metal ion selectivity or rely on a negative signal response (loss of fluorescence signal) as part of the detection process ^12, 14–16^.

PPIs underpin a huge number of biological processes critical to life ranging from gene transcription to energy production, to signal transduction ^17–21^, with PPI dysregulation contributing to many disease states ^20, 22^. Indeed, a protein is more likely to exist as part of a complex rather than act alone as a simple monomeric unit, with homodimers being the most commonly observed structural unit in the protein data bank (PDB) ^17, 18^. PPI interactions have been used as the basis for constructing FP-based sensors ^3, 23, 24^, including metal ions ^13^, through proximity-based processes such as Förster resonance energy transfer (FRET) ^25^ and fragment complementation (BiC) ^26^, These powerful and popular approaches have several drawbacks. One is the requirement for two different components, so restricting analysis to hetero-oligomerisation events thus missing out on frequently observed and equally important homo-oligomerisation.

The chromophore of the original FP, *Aequorea victoria* GFP (avGFP), exists in two forms: the neutral phenol state (CRO-OH) with an excitation maximum at ∼400 nm and the charged phenolate CRO-O^-^ form with an excitation maximum at ∼490 nm ^1, 27^. We have recently shown that the superfolder version of GFP (sfGFP) ^28^ can be engineered to change its spectral properties by switching the charged state on the chromophore by mutating H148 ^29^, a residue that stabilises the chromophore phenolate anion in the ground state^30–32^. When we replaced H148 by mutually reactive non-canonical amino acids, namely an azide group in one sfGFP and cyclo-alkyne in a second sfGFP, the CRO-OH state dominated in both ^33, 34^. When the two sfGFP “clicked” together to form a pseudo-homodimer there was a spectral shift with the CRO-O^-^ dominating and with the new dimeric form being brighter per monomer unit than sfGFP ^33^. While the basic premise of GFP monomer association provides a sensing mechanism, each sfGFP monomer unit should be chemically identical and not require genetic code reprogramming to incorporate a non-canonical amino acid.

Using only canonical amino acids we engineered sfGFP so that it’s spectral properties change on homodimerisation in a copper-dependent manner. By incorporating a cysteine at residue 148 together with a mutation that stabilises the dimer interface, we engineered sfGFP to switch to a highly fluorescent “on” dimeric state (Figure 1a). The dimer complex is 2.9 folder brighter than the starting wild-type monomeric sfGFP and has an up to 31 fold ratiometric switch. Structures of the monomer and dimeric states reveals a mechanism for switching based on a peptide bond flip and burial of critical water molecules central to GFP fluorescence. We also show that when coupled to naturally homodimerizing proteins, the copper-dependent kinetics of the engineered sfGFP increase, with the Cu^+^ being the preferred dimerisation promoting ion. A leucine zipper fused system allowed the newly engineered sfGFP to sense and respond to copper in bacterial cells.

**Figure 1.**
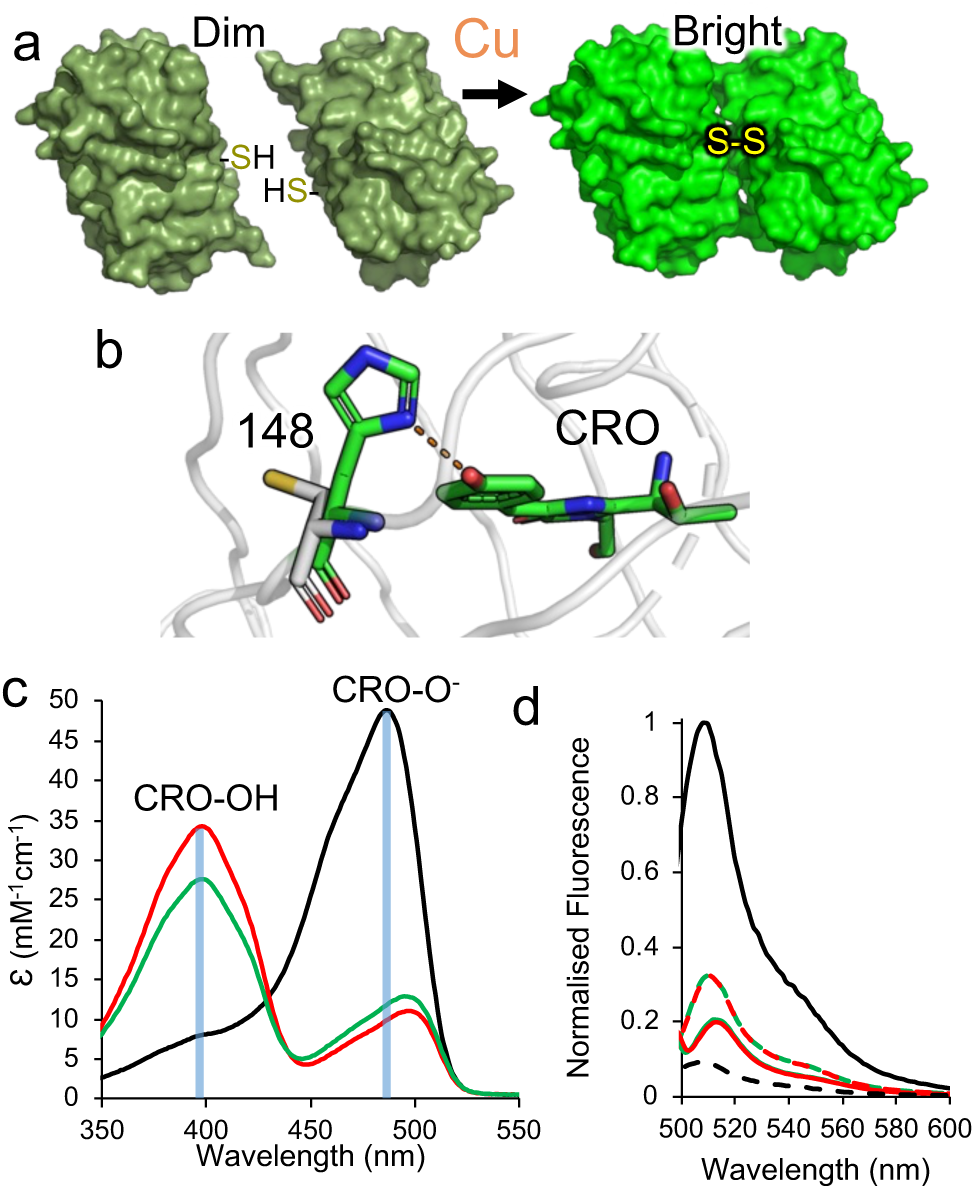
The effect of introducing cysteine residues at residue 148. (a) Overview of the homodimerization strategy. The starting monomers are dim with the CRO-OH form dominating but homodimerisation via a copper-dependent disulphide molecular bolt results in a brighter system through promotion of CRO-O^-^ state. (b) Modelling of the H148C mutation using short timescale molecular dynamics ^29^. The wt sfGFP is shown as green sticks with a H-bond between H148 and CRO shown as a dashed line. The H148C model is shown as grey sticks. Absorbance spectra (c) and fluorescence emission spectra (d) of wild-type sfGFP (H148, black), mGFP-diS1 (red) ^29^and mGFP-diS2 (green). In (d), the dashed line represent emission on excitation at 400 nm and full lines emission on excitation at 488-492 nm.

## Results and Discussion

### Switching the monomer chromophore protonation state

Modulating the charged state of the GFP chromophore phenol group provides a means to sense biological processes through changes in electronic excitation ^31–34^. In the starting wild-type sfGFP, the deprotonated CRO-O^-^ phenolate dominates over the neutral CRO-OH, with H148 capable of forming a H-bond with the chromophore phenolate group that stabilises the CRO-O^-^ form ^31–34^ (Figure 1b). Modelling previously suggested that mutating H148 to cysteine should remove the interaction with the chromophore ^29^; the weak Lewis acid behaviour of the cystine thiol group together with larger atomic radius of sulphur sterically blocks a conformation allowing H-bonding with the CRO and exposing the thiol group to the solvent (Figure 1b). Replacing H148 with cysteine generates the variant termed GFP-diS1, which remains monomeric after purification (Figure S1). Absorbance spectra shows that the major excitation peak is now 400 nm with a minor peak at 498 nm indicating a shift towards the CRO-OH state. The same is mirrored in fluorescence spectra with emission on excitation at 400 nm dominating over that on excitation at around 490 nm (Figure 1c and Table 1). Overall, the ratio of emission on excitation at wavelengths equivalent to the CRO-OH (∼400 nm) or CRO-O^-^ (∼490 nm) state (here on termed the Ex^CRO-O-^:Ex^CRO-OH^ ratio) switches from 7:1 in sfGFP to 1:4 in mGFP-diS1 (prefix m refers to monomeric).

**Table 1.** Spectral properties of sfGFP variants and dimers. [a] Excitation at 400 nm was negligible and so QY was not determined. [b] Value differs from that described in Pedelacq et al ^28^ but similar to those reported by Reddington et al ^31^ and Cranfill et al ^36^.

| Variant | $\lambda_{\max}$<br>(nm) | $\lambda_{\text{EM}}$<br>(nm) | $\epsilon$<br>(M<sup>-1</sup> cm<sup>-1</sup>) | $\Phi$<br>(%) | Brightness<br>(M<sup>-1</sup> cm<sup>-1</sup>) |
| --- | --- | --- | --- | --- | --- |
| **mGFP-diS1** | 400 | 509 | 34,000 | N/D | N/D |
|  | 498 | 513 | 11,100 | 50 | 5,550 |
| **dGFP-diS1** <sup>[a]</sup> | 400 | 509 | 11,400 | N/D | N/D |
|  | 488 | 509 | 142,600 | 64 | 91,264 |
| **mGFP-diS2** | 400 | 511 | 27,500 | N/D | N/D |
|  | 496 | 512 | 13,000 | 49 | 6,370 |
| **dGFP-diS2** <sup>[a]</sup> | 400 | 510 | 10,000 | N/D | N/D |
|  | 492 | 511 | 147,600 | 72 | 106,272 |
| **GFP-diS2-ZIP**<br>**(no Cu<sup>2+</sup>)** | 400 | 510 | 55,000 | N/D | N/D |
|  | 496 | 514 | 22,400 | 78 | 17,472 |
| **GFP-diS2-ZIP**<br>**(plus Cu<sup>2+</sup>)** | 400 | 510 | 17,700 | N/D | N/D |
|  | 493 | 511 | 188,400 | 85 | 160,140 |
| **sfGFP** <sup>WT [a,b]</sup> | 400 | 508 | 8000 | N/D | N/D |
|  | 485 | 511 | 49,000 | 75 | 36,750 |

### Copper-dependent dimerisation of GFP-diS1

Incubation of sfGFP-H148C mutants (termed GFP-diS1 from here on in) under aerobic conditions at high protein concentration (>50 μM) over a prolonged period (>5 hr) did not result in the formation of significant amounts of the dimer (Figure S1b); the absorbance spectra retained the characteristics shown in Figure 1c. This suggests that the cysteine thiol group cannot form the disulphide bond between individual GFP-diS1 monomers. It has previously been reported that Cu^2+^ can aid in the formation of disulphide bridges ^37^. On incubation with CuSO_4_, mGFP-diS1 dimerised to form dGFP-diS1 (prefix d refers to dimer), with ∼55% forming dimer after 1 hr (Figure S1b), similar to the yield observed for click-based dimerisation ^33^. The dGFP-diS1 dimer is stable and can be purified from the monomer by size exclusion chromatography (Figure S1c).

Dimerisation results in a shift in maximal absorbance to 488 nm (Figure 2a and Table 1), suggesting that CRO-O^-^ state now dominates dGFP-diS1. Moreover, molar absorbance for the CRO-O^-^ form increases ∼2.9 fold with a ∼2.5 fold increase in brightness compared to sfGFP (Figure 2a-b, Table 1). The greater than 2-fold enhancement in the spectral properties of dGFP-diS1 suggests a synergistic relationship on dimerisation where the dimer is greater than the sum of its parts, as observed previously for the click linked dimers ^33, 34^. Taking into account both the increase in emission on excitation at ∼488 nm and the decrease on excitation at ∼400 nm, we see a 31 fold signal change on dimerisation; the Ex^CRO-O-^:Ex^CRO-OH^ emission ratio at 510 nm changes from 0.58 for monomer to 18 for the dimer (Figure 2b).

**Figure 2.**
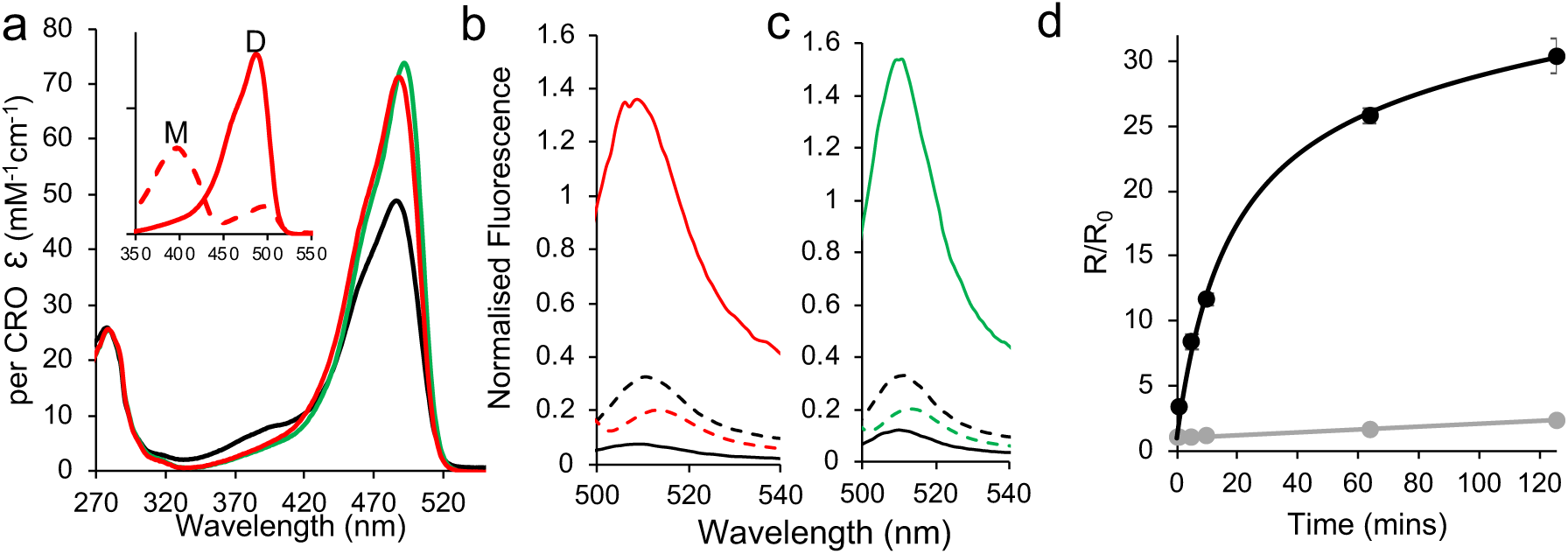
Dimerisation of sfGFP C148 variants. (a) Absorbance spectra of sfGFP^WT^ (black), dGFP-diS1 (red) and dGFP-diS2 (green). Inset is the change in absorbance on conversion of mGFP-S1 (dashed red line) to dGFP-diS1 (solid red line). The full absorbance spectra are shown in Figure S2. Emission spectra of (b) GFP-diS1 and (c) GFP-diS2, with red and green representing the dimers, respectively, and black the monomer. Solid and dashed lines represent excitation at either the optimal CRO-B wavelength (490-497nm) or CRO-A wavelength (400 nm), respectively. (d) Rate of dimerisation of GFP-diS1 (grey) and GFP-diS2. The data shown is for 10 μM of starting monomer with R/R_0_ representing the change in fluorescence emission ratio on excitation at either 388 nm or 483 nm.

The rate of mGFP-diS1 dimerisation is slow (Figure 2d), with an estimated second order rate constant of 0.1 nM/min. While formation of covalent complex means the system is non-equilibrium, an apparent *K*_D_ can be estimated to provide an insight into monomer affinity; for GFP-diS1, the relatively low level of association observed (Figure 2d) suggests that the *K*_D_ is >25 µM. It has been estimated that the dissociation constant for weak GFP dimerisation is 100 μM ^38^.

### Optimising the sfGFP interaction interface

The slow dimerisation kinetics of GFP-diS1 and low apparent affinity are likely due to interactions between the monomeric units being too transient and weak ^38^. Ideally, the kinetics should be faster and apparent *K*_D_ lower. Using the ncAA sfGFP dimer structure linked via residue 148 ^33^ (PDB 4nhn) as a guide, we aimed to optimise the GFP-diS1 interaction by targeting residue 206 that lies at the dimer interface (Figure S3) and has been shown previously to contribute to weak dimerisation of *Aequorea victoria* derived FPs ^36, 39^. We introduced the V206F mutation, so generating GFP-diS2. The GFP-diS2 variant is expressed and purified as a monomer (Figure S4a-b) and has similar spectral characteristics to mGFP-diS1 (Figure 1b-c and Table 1). The CRO-O^-^ absorbance peak at ∼490 nm has a slightly higher molar absorbance in mGFP-diS2 compared to mGFP-diS1 with a concomitant drop in ∼400 nm value. This could suggest GFP-diS2 improved the still weak tendency to self-associate so increasing the population of the CRO-O^-^ state (Figure 1b).

Again, Cu^2+^ is required to initiate dimerisation. In the presence of CuSO_4_ mGFP- diS2 dimerised with an efficiency of ∼80%, higher than that observed for mGFP-diS1, with the majority of dimerisation completed in the first 5 minutes (Figure S4b-c). Replacing CuSO_4_ with CuCl_2_ did not have any discernible change in the ability of GFP-diS2 to dimerise (Figure S4d). The switch in the major absorbance peak from 400 nm to 492 nm indicates that the CRO-O^-^ now dominates dGFP-diS2 (Figure 2a). This is also manifested in the fluorescence emission spectra with the higher emission on excitation at ∼490 nm (Figure 2c). As with dGFP-diS1, dGFP-diS2 was brighter (2.9 fold) than wild-type sfGFP (Table 1). Indeed, dGFP-diS2 is brighter than dGFP-diS1 due to slightly higher molar absorbance and quantum yield (Table 1). The Ex^CRO-O-^:Ex^CRO-OH^ (400 nm:492 nm) ratio on measuring emission at 510 nm changes from 0.56 to 13.4, which equates to a 24 fold change signal on dimerisation. Significantly, there is the large increase in dimerisation rate (Figure 2d), with the second order rate constant increasing two orders of magnitude to 11 ± 1 nM^-1^min^-1^. The apparent *K*_D_ is estimated to be 4.6 ± 0.2 μM, equivalent to a moderate protein-protein interaction. The dimerisation process is reversible as dGFP-diS2 can be converted back into its monomer components with monomeric spectral characteristics using DTT (Figure S5). Thus, GFP-diS2 is a distinct improvement on GFP-diS1 in terms of its copper-dependent homodimerisation properties.

### Structural basis for fluorescence switching in dimerisation

To fully understand the basis behind dimerization-dependent fluorescence switching, we determined the structure of the GFP-diS2 monomer and dimer (see Table S1 for structural statistics). In mGFP-diS2, H148C has a major impact on the local organisation of the structure that directly impacts on the interactions with the CRO (Figure 3a-b and Figure S6). The backbone of residues 147-149 is twisted compared to sfGFP^WT^ resulting in a register shift of S147 and C148 (Figure 3b and Figure S6). The change in position of C148 results in the loss of the H-bond to the CRO and becomes solvent exposed making it amenable to forming a disulphide bond. S147 now points towards the chromophore with the OH groups within H-bonding distance of each other (Figure 4b). The new potential H-bond between CRO and S147 does not however appear to compensate for the H148C mutation in terms of promoting the phenolate chromophore form. Indeed, compared to wt sfGFP, the water network is less extensive around the phenol group for mGFP-diS2, with the sole water present potentially undergoing dynamic exchange with the bulk solvent via the channel shown in Figure 3b. Such water networks are known to play a key chromophore charge state ^33, 40, 41^. The side chain orientation of E222 and thus its interaction with S205, a critical interaction involved in proton shuttling that promotes the CRO-O^-^ form ^42, 43^, is also altered in mGFP-diS2; as a result, the distance between E222 and S203 increased to 3.7Å (from 3.0Å in sfGFP). Thus, the cumulation of these structural changes are likely to promote CRO-OH on the introduction of the H148C mutation. The V206F mutation does not have any significant impact on the local structure of sfGFP with both backbone and side-chain configurations of the mutated residue and surrounding residues being similar (Figure S7).

**Figure 3.**
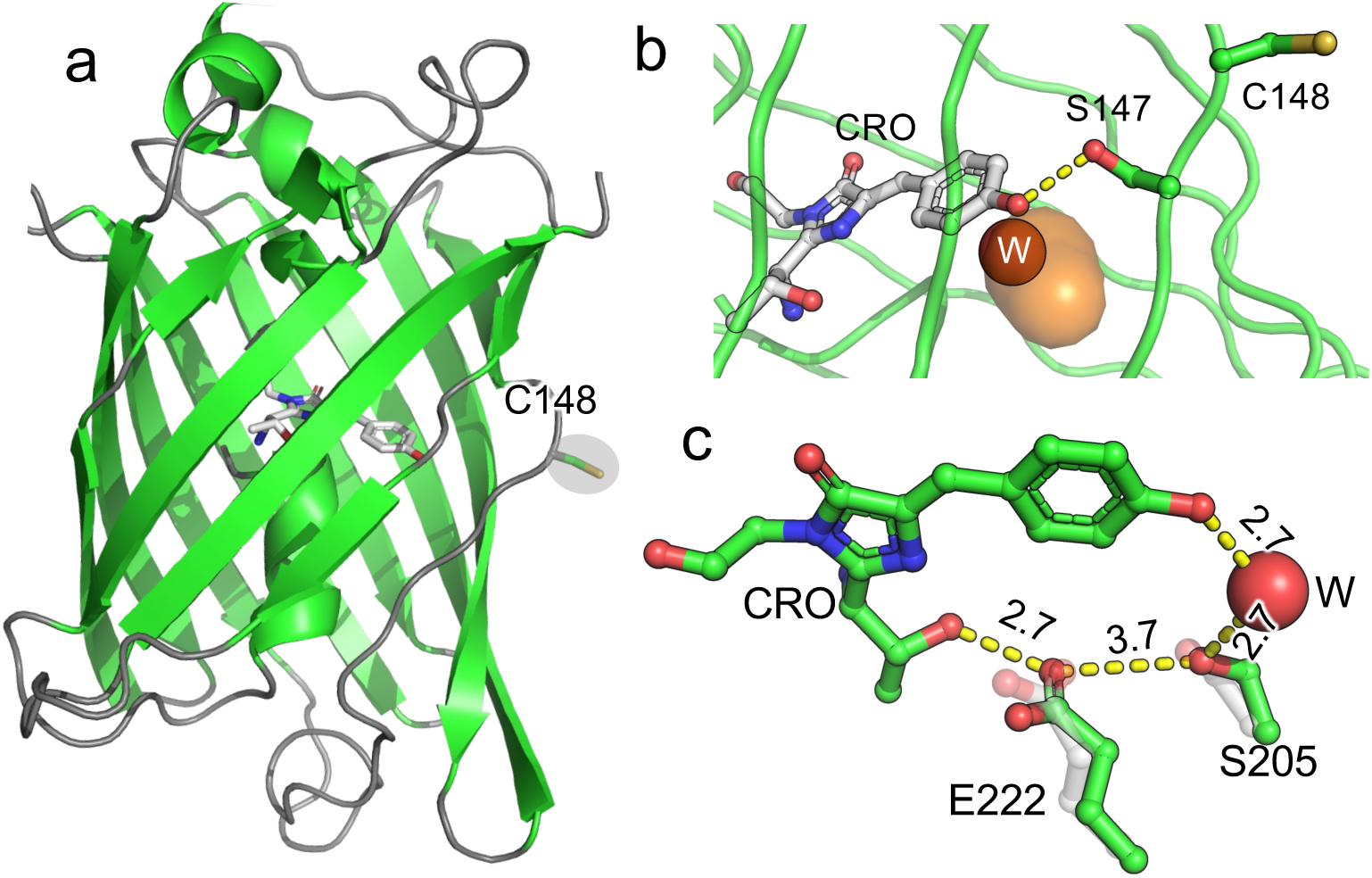
Structure of mGFP-diS2 (PDB 8c1x). (a) Overall structure mGFP-diS2 with the chromophore shown as grey sticks and C148 shown as sticks and highlighted. (b) Structure of the chromophore and residues 147-148. Dashed lines represent potential H-bonds with the phenolate and either S147. A channel through to the chromophore (CRO) (coloured orange) is shown together with a single conserved water molecule (W, red sphere) within the channel. The channel was calculated using CAVER3.0 with a probe size of 1.0 Å. (c) Residues involved in the classical proton transfer network related to promotion of the phenolate chromophore form. The grey residues are the equivalent position of S205 and E222 in sfGFP. Distances shown are for mGFP-diS2 and are in Ångstroms.

**Figure 4.**
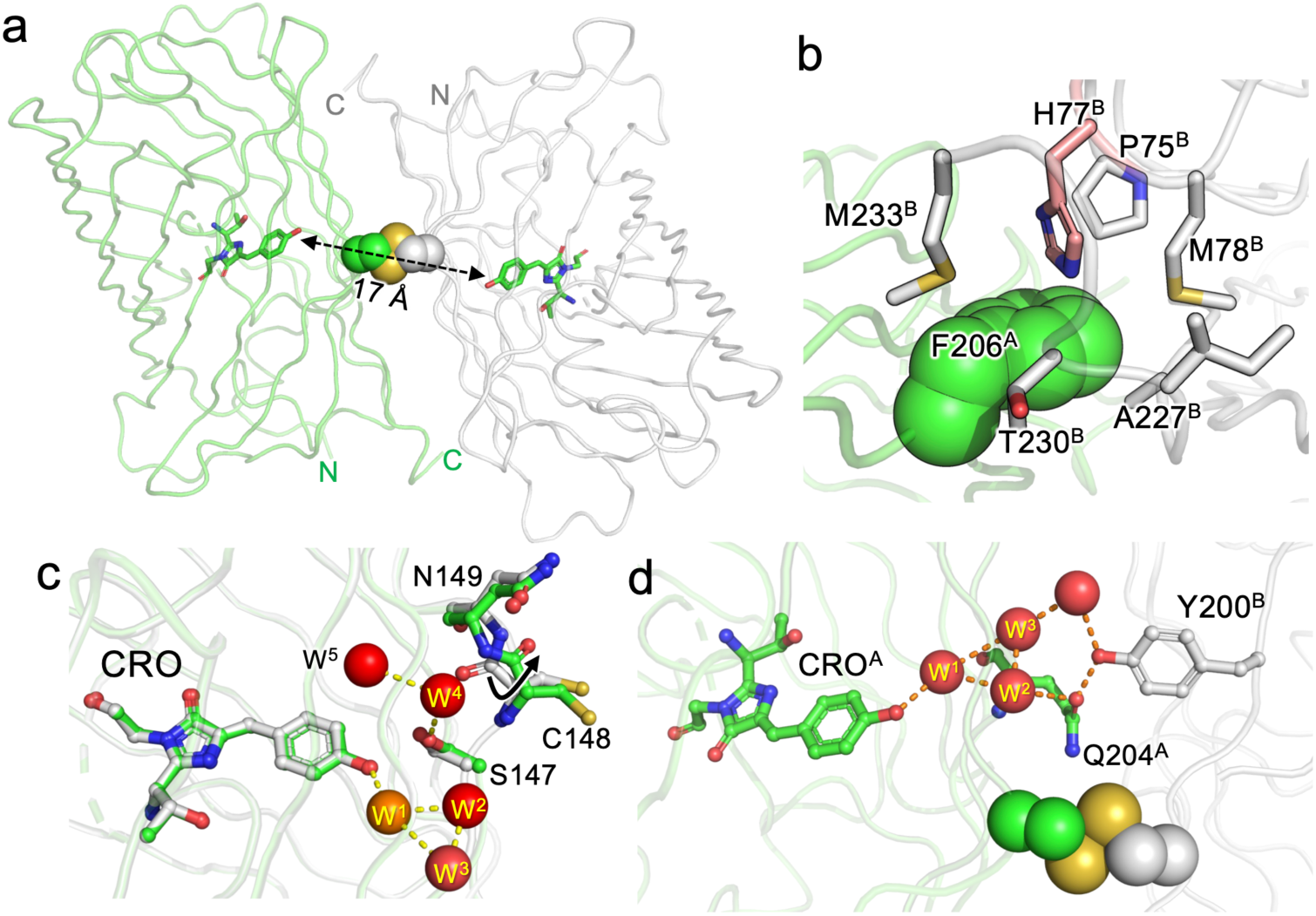
Structure of dGFP-diS2. (a) Overall structure of dGFP-diS2 showing one monomer (green) in relation to the other (grey). The chromophores are shown as sticks and coloured green and the disulphide bridge is shown as spheres. The electron density of the disulphide bridge is shown in Figure S8. The distance between the two chromophores is shown by the dashed double arrowhead line. (b) The role of the V206F mutation in stabilising the dimer interface. Residues with superscript A come from one subunit and are coloured green and those with superscript B are from the second subunit and coloured grey. The H77^B^ is highlighted as part of a pi-cation interaction. (c) The local chromophore environment with water molecules shown as red spheres. The monomeric mGFP-diS2 is overlaid and coloured grey with the conserved W^1^ water molecule present in both structures coloured as an orange sphere. (d) Inter-subunit network between CRO^A^ and Y200^B^.

The overall structure of dGFP-diS2 is shown in Figure 4a. The two dimers have a defined, symmetrical interface comprising the same residues from each monomer unit covering 910 Å^2^ (as defined by PISA^44^). The residues that contribute dimer interface are detailed in Tables S2 and S3. The disulphide bridge has two potential configurations (see Figure S8 for electron density) and is exposed to the solvent explaining the ability of DTT to break the dimer apart (Figure S6). The introduction of the V206F mutation plays an important role in promoting the dimer formation as it makes interactions with 6 residues in the neighbouring monomer, including a pi-cation interaction with H77 (Figure 5b).

**Figure 5.**
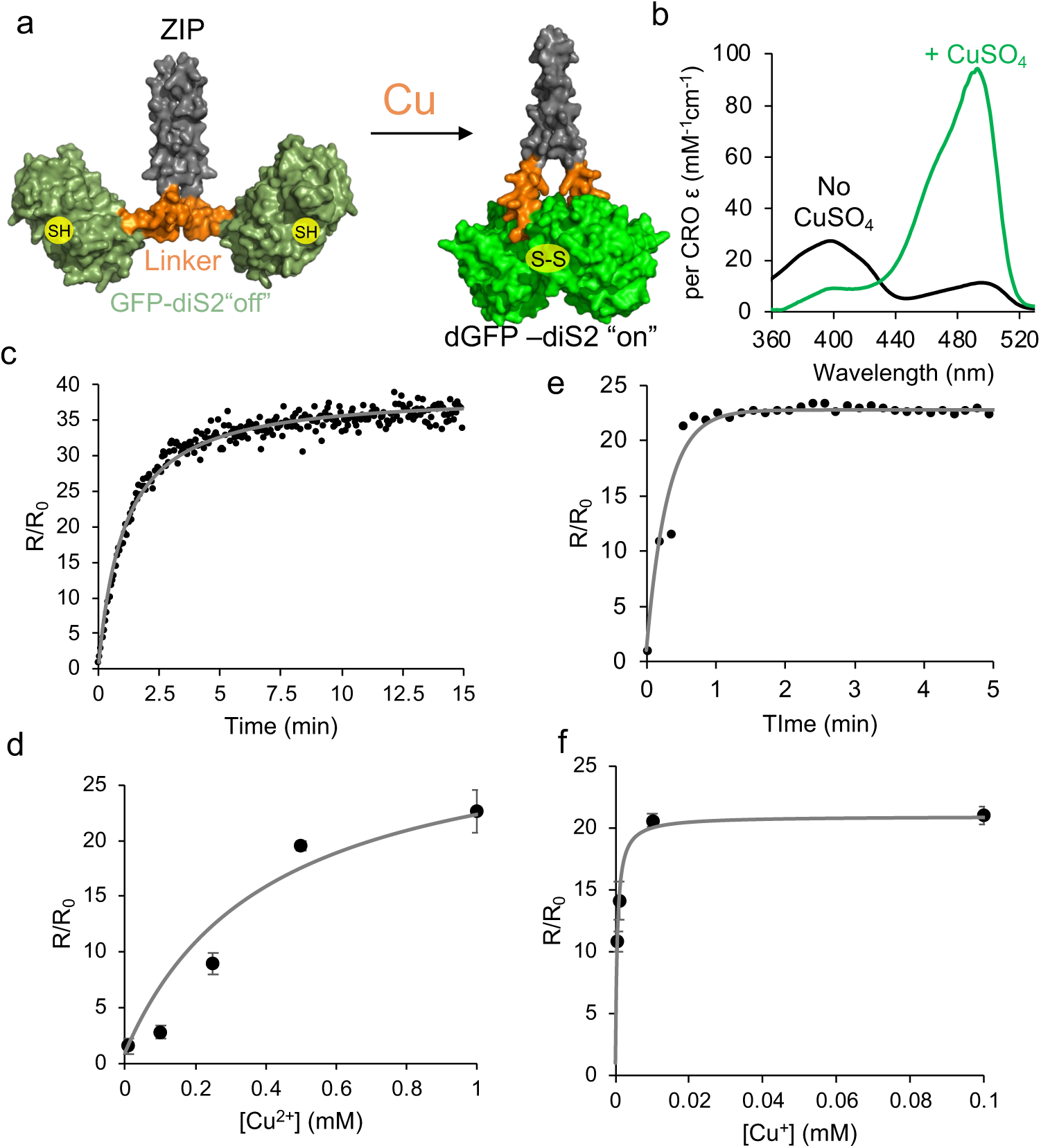
Dimerisation of GFP-diS2 fused to a leucine zipper element. (a) Schematic representation of activation mechanism. The leucine zipper (ZIP) is coloured grey, linker sequence orange and the GFP-diS2 component green. (b) Absorbance spectra of GFP-diS2-ZIP in the absence (black) and presence (green) of 1 mM CuSO_4_. *In vitro* association kinetics of GFP-diS2-ZIP in the presence of (c) Cu^2+^ (1 mM CuSO_4_) or (e) Cu^+^ (250 μM CuSO_4_ plus 1 mM ascorbate). The influence of either (d) Cu^2+^ or (f) Cu^+^ on the GFP-diS2-ZIP fluorescence ratio.

The likely mechanism of CRO state switching is changes in the local chromophore environment on dimerisation. Monomeric and dimeric GFP-diS2 are very similar with a Cα RSMD of 0.119 Å and local side chains around the chromophore occupying similar positions (Figure S9a). The main changes are increased solvation of the chromophore and a plane flip in the position of the C148-N149 peptide bond (Figure 4c and S9b). Peptide bond plane flipping is relatively rare and may be the rate-limiting step in the dimerisation processes^45, 46^; it could also provide a rationale for the requirement for Cu^2+^ to promote dimerisation. The water molecule close to the CRO (W^1^) observed in mGFP-diS2 (Figure 3b) is still present in the dimer but 4 additional water molecules are now observed. Together with W^1^, the W^2^ and W^3^ water molecules form a H-bond network to the CRO with these waters being planer to the CRO (Figure 4c). W^4^ and W^5^ are above the plane of the CRO and occupy the space available due to rearrangement of the C148-N149 peptide bond; they form a H-bond network with S147. In the broader picture of the dimer, waters W^1^, W^2^ and W^3^ form a long-range polar network that spans the dimer interface incorporating Y200 from the second protomer (Figure 4d). The formation of organised water networks in avGFP-like proteins close to the CRO has been postulated previously to promote the formation of the ground state CRO-O^-^ form ^32–34^. The water networks observed here (Figure 4c-d) are likely to play a similar role by facilitating proton abstraction which can then be shuttled though the neighbouring water network at the dimer interface. These local water molecules together with S147 can help stabilise the negative charge so promoting CRO-O^-^. Crucially, these water molecules are also largely buried at the dimer interface so restricting their exchange with bulk solvent (Figure S10). Recent work has shown that the structurally conserved water molecule W^1^ rapidly exchanges with the bulk solvent in sfGFP^29^. The potential to restrict exchange of water molecules may contribute to the improved spectral properties on dimerisation both here and elsewhere ^33, 34^.

### Fusing GFP-diS2 to a homo-dimerising component

The switch from CRO-OH to CRO-O^-^ provides a means to monitor copper levels and also protein dimerisation through ratiometric changes in fluorescence emission on excitation at two separate wavelengths (∼400 nm and ∼490 nm). The N- and C-termini are on the same face (Figure S11), so should not preclude association if fused to naturally homo-dimeric proteins. To assess the effect of fusing GFP-diS2 natural homodimerizing protein component, we employed the leucine zipper region (ZIP) of transcription factor GCN4 ^47^. In this scenario, the leucine zipper region should result in GFP-diS2 becoming an obligate homodimer (Figure 5a). Addition of copper ion should result in rapid switching “on” of fluorescence emission on excitation at ∼490 nm due to the localisation of the two GFP-diS2 components.

In the absence of CuSO_4_, the homodimer construct now termed GFP-diS2-ZIP, exhibited spectral characteristics similar to mGFP-diS2 (Figure 5b and Table 1), with the ∼400 nm absorbance peak equivalent to CRO-OH dominating. On addition of CuSO_4_, GFP-diS2-ZIP quickly switches to the CRO-O^-^ as indicated by the dominant absorbance at ∼493 nm (Figure 5b). As with the monomer-to-dimer conversion of GFP-diS2, the chromophore molar absorbance coefficient was enhanced compared to sfGFP and dGFP-diS2 (Table 1). The Ex^CRO-O-^:Ex^CRO-OH^ fluorescence emission ratio was also higher in the obligate homodimer, with a 37 fold difference compared to 27 fold change when forming dGFP-diS2. The rate of conversion was also far quicker (Figure 5c) with a second order rate constant of 810 ± 30 nM^-1^min^-1^, ∼65 fold quicker than monomer-to-dimer conversion; for a 1 μM sample of GFP-diS2-ZIP conversion was essentially complete within 5 min (Figure 5c). Thus, coupling GFP- diS2 to a known homodimerisation element results in a much improved GFP association and thus switch in fluorescence. The change in emission ratio is also dependent on Cu^2+^ concentration suggesting GFP-diS2-ZIP can be used as sensor cellular copper sensor (Figure 5d). At Cu^2+^ concentrations <=10 μM, the rate of switching is slow (Figure S12a) with the end point change being relatively small (Figure 5d). The ratiometric switch also appears to be specific to copper ions as other biologically relevant metal ions did not change the Ex^CRO-O-^:Ex^CRO-OH^ emission ratio (Supporting Table S4).

Inside the reducing environment of the cell, Cu^2+^ is reduced to Cu^+^ and is mainly transported into the cell as the Cu^+^ form ^8,10^. For use inside the cell, we assessed the impact of reducing conditions and Cu^+^ on GFP-diS2-ZIP fluorescence switching. Cu^+^ is commonly produced in the presence of ascorbate, a natural biological reducing agent ^48^. The rate of association increased considerably in the presence of Cu^+^, with fluorescence switching occurring in <1 min (Figure 5e), with an estimated second order rate constant of 3118 ± 465 nM^-1^ min^-1^. Cu^+^ also display a concentration dependence in fluorescence switching (Figure 5f and Figure S12b), as with Cu^2+^. Ascorbate itself has little effect on the spectral properties of GFP-diS2-ZIP (Figure S13). It is not clear why Cu^+^, a reducing agent, should promote bringing the two GFP-diS2 units close to each and instigate switching using the proposed disulphide mechanism above. Cu^+^ does have high affinity for thiolates capable of forming a bis-thiolate complex (e.g. in CSP1 ^49^). Here, Cu^+^ could be acting as the bridge linking together the GFP-diS2 units via the engineered cysteine residues.

We then tested the GFP-diS2-ZIP system in situ by monitoring changes in fluorescence ratio in *E. coli*. The R_0_ ratio (Ex 390 nm:Ex 490 nm at 0 s) ratio was ∼0.60, close to the in vitro value of 0.58 reported above. This suggests that free copper ion is very low in ambient conditions, which is in line with previous work ^8,50^. On addition of CuSO_4_ to the cell suspensions, there is an initial lag of ∼500 s, followed by an increase in fluorescence ratio; a plateau is reached after ∼55 min (Figure 6a). The change in the maximal fluorescence ratio is 7.2 fold, which equates a sub μM Cu^+^ concentration and/or 100 μM Cu^2+^, based on the metal ion dependent concentration curves in Figure 5.

**Figure 6.**
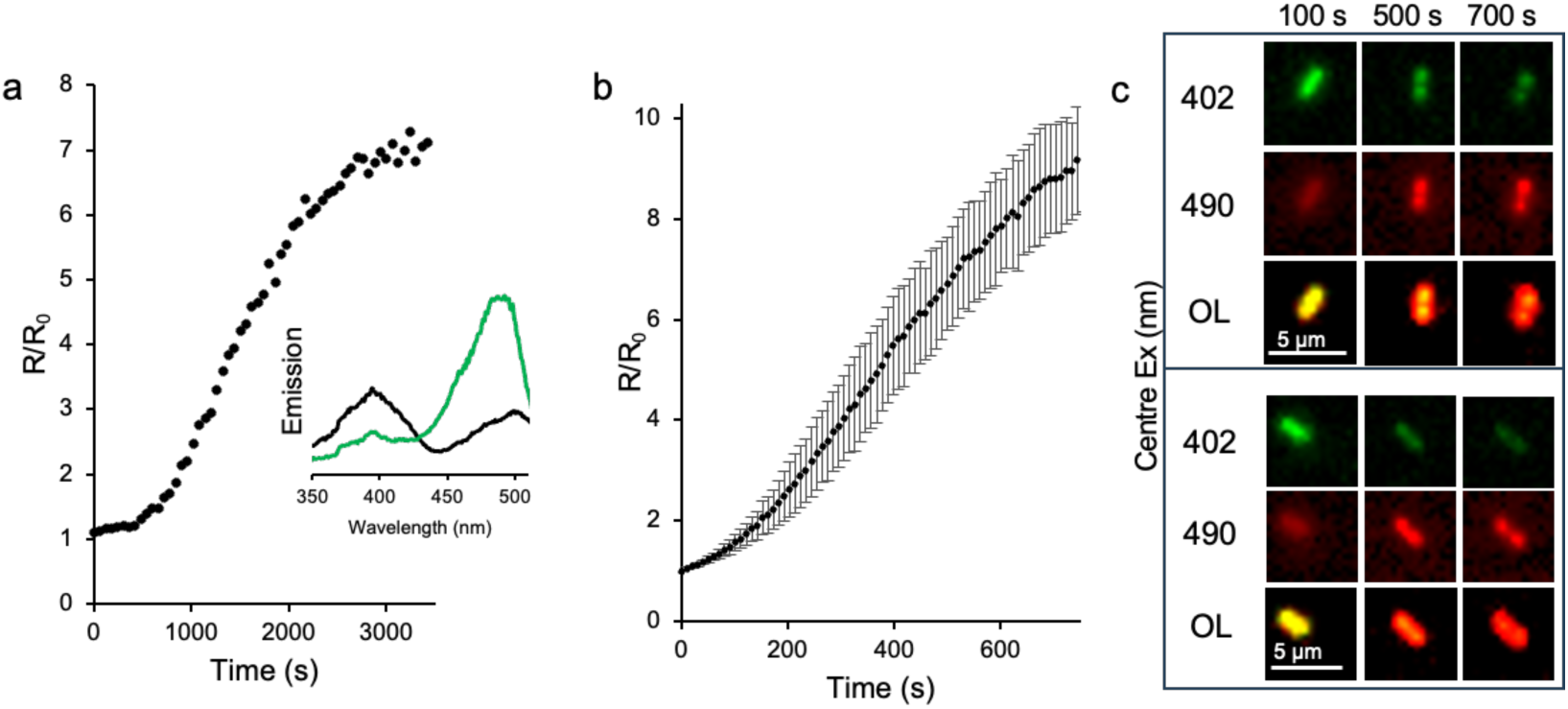
*In situ* response of GFP-diS2-ZIP to copper. (a) Change in fluorescence emission ratio (Ex 390 nm: Ex 490 nm) of bulk *E. coli* cell culture on addition of 1 mM CuSO_4_. Inset is the fluorescence emission spectra (emission at 520 nm) measure before (black) and 3600 s after (green) the addition of 1 mM CuSO_4_. (b) Change in fluorescence emission ratio (Ex 405 nm: Ex 470 nm) of individual *E. coli* cells on additional of 1 mM CuSO_4_. A total of 23 different cells were analysed from widefield imaging. The error bars represent the standard deviation between the 23 different cells. (c) Representative widefield images at the different centre excitation wavelength of individual *E. coli* cells at different time periods. OL refers to the overlay of the two individual excitation wavelengths. The accompanying movies over the full time course can be found as Supplementary Movies 1 and 2.

GFP-diS2-ZIP can also be used as a cell imaging probe. Fluorescence microscopy allows the impact of copper on individual cells to be investigated. As with the bulk cell culture, an initial lag is observed of ∼60 s (plus 4-5 min preparation before imaging) after adding CuSO_4_ to the cells. After the initial lag phase, the fluorescence emission ratio increases until it reaches an R/R_0_ plateau at around 9. This equates to a sub micromolar Cu^+^ concentration, the form of copper that is thought to predominate inside the cell under reducing conditions ^11^. The cells analysed appear to retain their general cellular morphology with the changes in fluorescence levels clearly evident (Figure 6c).

Our data suggests that copper can enter *E. coli* with available levels likely to be in the 10-100s nM range (10^-8^-10^-7^ M). Copper import into *E. coli* is largely unknown but the initial lag period suggests there is a requirement to induce production of import systems such as the recently report AZY operon system ^51^. It was originally thought that only efflux systems were induced in response to high copper concentrations ^8,10, 11^. Cellular copper is thought to be ∼10 μM but the vast majority is bound to proteins leaving the available intracellular free copper ion at <10^-18^ M or just a few atoms per cell ^52^; this can rise to 10^-15^ M when overloaded, as is likely the case here when a high (1 mM) CuSO_4_ concentration is added to cells. The reported overload level is still much lower than our data here suggests. What is clear that our GFP-diS2-ZIP system can respond dynamically to changes in cellular copper concentrations generating a positive and large (compared to existing genetically encoded copper sensors) signal change.

## Conclusion

We have engineered a well-utilised GFP to change it spectral characteristics in response to copper-dependent homo-dimerisation. The molecular basis of action suggests that organised water molecules play a key role in promoting proton transfer from the chromophore causing a switch in its charged ground state from neutral phenol to negatively charged phenolate. This in turn changes the primary excitation wavelength providing a measurable response on homo-dimerisation. The requirement for CuSO_4_ provides a means to measure copper levels without interference from other metal ions and to our knowledge is the first genetically encoded ratiometric copper sensor with a large positive gain of signal. There is the potential to transfer the system to monitor more generally homodimerisation, arguably the most common of all PPIs in biology, by replacing the leucine zipper component with another homodimerisation element. The need for copper and the disulphide bridge is currently a limiting aspect but further engineering to optimise the dimer interface and promote the 148-149 peptide bond plane flipping may potentially result in a non-covalent system that can monitor reversible and dynamic exchange PPI events.

## Methods and Materials

### Protein engineering and purification

The gene encoding sfGFP was resident in the pBAD plasmid (arabinose inducible) ^31^ or the pCA24N P50-GFP fusion (see Supporting Information for gene fusion construction) were used as the template for whole plasmid inverse Q5 DNA polymerase mutagenesis procedure (NE Biolabs). The H148C and V206F mutations were introduced using the following primer pairs: H148C, 5’- GCGGTAATATACACATT**<u>GCA</u>**GCTGTTG-3’ and 5’-

CGATAAACAGAAAAATGGCATCAAAGCG-3’; V206F, 5’-

GAGCAAAGATCCGAATGAAAAACGTG-3’ and 5’-

AG**<u>AAA</u>**GCTCTGGGTGCTCAGATA-3’. All mutations were confirmed by DNA sequencing (Eurofins Genomics). The GFP-diS2-ZIP construct was synthesised by Twist Bioscience and cloned into the pET28a (IPTG inducible). Proteins were purified essentially as described previously ^31, 53^ and with details provided in the Supporting Information.

### Dimerisation of sfGFP variants

CuSO_4_ (Sigma-Aldrich) was added to sfGFP variant protein samples to aid dimerisation through formation of a disulphide bond ^54^. Dimerisation of the sfGFP variants in 50 mM TrisHCl, pH 8, was initiated by the addition of CuSO_4_ to a final concentration of 1 mM and then left to incubate for varying amounts of time. For association kinetics, protein concentrations ranged from 0.1μM to 0.5 μM with samples taken periodically for analysis. The dimerisation of sfGFP variants used in this study was a second order rate reaction, as homodimerisation is recorded as A +A → A_2_ ^55^. The second order rate constant was calculated using the equation v = *k*[A]^2^ where v is the rate, *k* is the rate constant (μM^-1^sec^-1^) and A is the monomer protein concentration (μM). To calculate the percentage of sfGFP that has dimerised, the emission ratios on excitation of the CRO-OH (390-400 nm) or CRO-O^-^ (485-495 nm) were recorded at various time points as described below. For preparation of large quantities of dimer, samples of up to 50 mg/mL of protein were incubated with 1 mM CuSO_4_ for 1 hr with mixing. The monomer and dimer were separated by SEC as described in the Supporting Information.

### Absorbance and fluorescence spectroscopy

UV-visible (UV-vis) absorption spectra were recorded on a Cary 60 spectrophotometer (Agilent) in a 1 cm pathlength quartz cuvette (Hellma). Spectra were recorded from 200 – 600 nm at a rate of 300 nm/min. Extinction coefficients were calculated using the Beer-Lambert law by using known concentration of protein as determined by the DC protein assay (BioRAD) with sfGFP as the calibration standard and verified by comparison with sfGFP’s 280 nm absorbance (sfGFP, ɛ_280_ = 25,400 M^-1^cm^-1^). Steady state emission and excitation fluorescence spectra were measured using Varian Cary Eclipse Fluorimeter and a 10 mm x 2 mm QS quartz cuvette (Hellma). Spectra were recorded with 5 nm excitation/emission, slit width at a rate of 300 nm/min. Emission spectra were recorded at a fixed excitation wavelength according to the excitation maximum of the variant. Excitation spectra were recorded at the fixed wavelength according to the maximum emission. Unless otherwise stated, spectral scans were recorded for at 0.5 μM in 50 mM Tris-HCl, pH 8.0, for monomers and at 0.25 μM for dimers. For GFP-diS1 and GFPdiS2 dimerisation kinetics, a CLARIOstar Plus plate reader (BMG LabTech) was also used. The bandwidth of filters used was 8 nm. As the minimum distance allowed between filters is 30 nm, the emision was recorded at 516 upon excitation at 388 nm and 483 nm. Each measurement was performed in triplicate using a NuncTM F96 MicrowellTM black polystyrene plate (Thermo fisher). To calculate Ex^CRO-O-^:EX^CRO-OH^ ratios, emission values upon excitation at major absorbance peak around 490 nm were divided by emission values when recorded at ∼400 nm. Extact wavelengths are stated in the main text. The ratio are presented as R/R_0_ where R is the ratio at a particular time point and R_0_ the ratio at the start of the measurement. Quantum yield was determined using a fluorescein standard as described previously ^32^.

### Structure determination

Purified proteins samples in 50 mM Tris-HCl, pH 8 at ∼10 mg/mL were screen for crystal formation using sitting drop vapour diffusion across a wide variety of conditions as described by the PACT *premier*TM and JCSG-*plus*TM HT-96 broad crystallisation screens (Molecular Dimensions). Crystallisation plates were stored at 23°C and crystal growth monitored. Crystals were harvested, 1 mM ethylene glycol added and flash frozen in liquid nitrogen. X-ray diffraction data was obtained at Diamond Light Source, Harwell, UK on beamline I03. Crystals of mGFP-diS2 were in the P 1 2_1_ 2 space group grew in condition G12 on the PACT Premier screen (0.2 M Sodium malonate dibasic monohydrate, 0.1 M Bis-Tris propane, pH 7.5, 20 % w/v PEG 3350). The crystals diffracted to the highest resolution of 1.89 Å. Crystals of dGFP-diS2 were in the P 2 2 2_1_ space group and grew in condition G7 on the PACT Premier screen (0.2 M sodium acetate trihydrate, 0.1 M Bis-tris propane, pH 7.5, 20% w/v PEG 3350). The crystals diffracted to the highest resolution of 1.79 Å. The structures were determined by molecular replacement, with the sfGFP structure (PDB 2b3p^28^) as the reference, essentially as described previously ^32, 33^. Full diffraction data and refinement statistics can be found in Table S1. The PDB entry for mGFP-diS2 is 8c1x and is 8bxp for dGFP-diS2.

### E. coli cell imaging

Bacteria were induced with 1 mM IPTG and grown to mid log phase before being concentrated by centrifugation tenfold to increase the number of bacteria present within an individual field of view. Cells were mixed with CuSO_4_ (final concentration of 1 mM), before 20 uL were placed on a fresh microscope slide and covered with a 25mM #1.5 glass coverslip and immediately placed on the microscopy for imaging. Wide-field epi fluorescence measurements were conducted on an inverted Olympus IX73 microscope and a Prior Lumen200Pro light source using filter set 89000, (Chroma, Vermont, U.S.A.) selecting the ET402 nm/15 nm or ET490 nm/20 nm as the excitation filter and the ET525 nm/36 nm as the emission filter. The fluorescence emission was detected with a Hamamatsu ORCA-flash 4.0 V2 sCMOS Camera operated utilizing the HCImage software package (Hamamatsu). An Olympus UPlanSApo 20× oil immersion objective with an NA of 0.75 was used to collect sequential images at full resolution (2048x2048 pixels at 1xbinning) with exposure times of 500 ms for each channel, with a 10 second interval over 75 timepoints. To reduce any impact of sample drift on quantification, cell images were registered in xy using the “Linear stack alignment with SIFT multichannel” plugin ^56^ in FIJI ^57^. A total of 23 individual bacteria were selected through the use of a 5 pixel diameter circular region of interest, and the intensity of each channel within each region of interest (ROI) was measured. Background measurements were calculated from the average of 5 ROI containing no bacteria at time 0, and subtracted from each measured timepoint. Finally, a ratio was determined between centred 402 and 490 nm excitation and the R/R_0_ calculated for each individual bacteria, before being averaged across all bacteria and plotted against time.

## Supporting information

SuppInfo

## Author contributions

All authors contributed to the writing of the paper and analysing data. RDA designed monomer variants, prepared in the monomer variants, analysed the initial monomer variants (including dimerisation potential) and determined their structure. DV designed and generated the LZ variants, analysed the LZ variants and undertook the in cell analysis. KDH contributed to the LZ variant analysis and cell imaging. KA contributed towards LZ variant analysis and cell imaging. PJR collected structural data and helped with structure determination and refinement. PDW contributed to cell imaging and data analysis. DDJ conceived and directed the project, contributed to variant design and data analysis.

## Acknowledgement

We would like to thank the staff at the Diamond Light Source (Harwell, UK) for the supply of facilities and beam time, especially beamline I03 staff, under beamtime code mx18812. DDJ would like to thank the following funders: KESS2 (Knowledge Economy Skills Scholarship European Regional Development Fund via Welsh Government) studentship in partnership with Tenovus (to RDA), a Royal Thai Embassy studentship to DV, Tabuk University and the Royal Embassy of Saudi Arabia (to KA), BBSRC (BB/Z514913/1), and EPSRC (EP/V048147/1). The authors would like to thank the Cardiff School of Biosciences Protein Technology Hub for helping with the production and analysis of proteins, and the Cardiff University Bioimaging Hub Core Facility (RRID: SCR_022556) for their support and assistance in this work.

