## Supplementary material for "Chromophore charge-state switching through copper-dependent homodimerisation of an engineered green fluorescent protein": SuppInfo

**Supplementary Information.**

**Supporting Methods**

**Sequence of the GFP-diS2-ZIP resident in pET28a.**

ATGGTTAGCAAAGGTGAAGAACTGTTTACCGGCGTTGTGCCGATTCTGGTGGA
ACTGGATGGTGATGTGAATGGCCATAAATTTAGCGTTCGTGGCGAAGGCGAAG
GTGATGCGACCAACGGTAAACTGACCCTGAAATTTATTTGCACCACCGGTAAAC
TGCCGGTTCCGTGGCCGACCCTGGTGACCACCCTGACCTATGGCGTTCAGTGC
TTTAGCCGCTATCCGGATCATATGAAACGCCATGATTTCTTTAAAAGCGCGATG
CCGGAAGGCTATGTGCAGGAACGTACCATTAGCTTCAAAGATGATGGCACCTAT
AAAACCCGTGCGGAAGTTAAATTTGAAGGCGATACCCTGGTGAACCGCATTGAA
CTGAAAGGTATTGATTTTAAAGAAGATGGCAACATTCTGGGTCATAAACTGGAAT
ATAATTTCAACAGCTGCAATGTGTATATTACCGCCGATAAACAGAAAAATGGCAT
CAAAGCGAACTTTAAAATCCGTCACAACGTGGAAGATGGTAGCGTGCAGCTGG
CGGATCATTATCAGCAGAATACCCCGATTGGTGATGGCCCGGTGCTGCTGCCG
GATAATCATTATCTGAGCACCCAGAGCTTTCTGAGCAAAGATCCGAATGAAAAA
CGTGATCATATGGTGCTGCTGGAATTTGTTACCGCCGCGGGCATTACCCACGG
TATGGATGAACTGTATAAAGGCAGCACCAAAGATAAAGAAAACCTGTATTTTCA
GTCTAAGCAGCTTGAGGACAAAGTCGAGGAACTGCTATCCAAGAACTATCATCT
GGAGAACGAGGTGGCTCGTCTGAAGAAACTGGTGGGTGGTTCTCACCATCATC
ATCACCATTAA

**Protein purification.**

The wild-type sfGFP and it's variants in the pBAD vector (Amp[R]) were transformed into *E. coli* Top 10[TM] (Invitrogen, Paisley, UK). The pET28a GFP-diS2-ZIP (Kan[R]) and the pCA24N P50-GFP (Cm[R]) variants were transformed into *E. coli* BL21 (DE3) (NE Biolabs) for recombinant protein expression. A single colony was taken from the transformation and used to inoculate a 10 mL 2xYT starter culture supplemented with suitabble selective antibiotic. The starter cultures were placed in a shaking incubator overnight at 37 °C. For the pBAD-based variants, the starter culture was used to inoculate 1L culture of autoinduction media supplemented with ampicillin (50 µg/mL) [1]. The autoinduction cultures were incubated for a further 18-24 hr at 37°C. For the pCA24N P50-GFP variants, 1L of 2xYT supplemented with chloramphenicol (35 µg/mL) was inoculated with the starter culture and incubated at 37 °C with shaking (200 RPM). At an $OD_{600}$ of ~0.8, IPTG was added to a final concentration of 0.4 mM and incubated overnight with shaking at 28 °C. The cells were then pelleted by centrifugation and then resuspended 50 mM Tris-HCl, pH 8.0. Cells were lysed using the French Pressure Cell. Soluble cell lysate was then separated from insoluble fractions via centrifugation at 25,000*g* for 40 mins. Clarified cell lysate was passed through a 5 mL His Trap™ HP column (Cytiva) equilibrated in 50 mM Tris-HCl, pH 8.0 buffer containing 10 mM imidazole. Bound target protein was then eluted by the addition of the 50 mM Tris-HCl, pH 8.0 buffer containing imidazole at a gradient from 10 to 500 mM. Samples of each fraction were taken and used to check for purity via SDS-PAGE. When required size exclusion chromatography (SEC) was performed to further purify samples and to analyse the oligomeric state. SEC was performed with either HiLoad™ 16/600 Superdex™ S75 pg (Cytiva) or HiLoad™ 16/600 Superdex™ S200 pg (Cytiva).

**Table S1.** Crystallographic statistics from x-ray diffraction refinement and final bond parameters for the GFP-diS2 crystal structures

| | **mGFP-diS2** | **dGFP-diS2** |
|---|---|---|
| **Data collection/reduction statistics** | | |
| PDB ID | 8c1x | 8bxp |
| Wavelength (Å) | 0.81532 | 0.81532 |
| Beamline (DLS, UK) | I03 | I03 |
| Space group | P 1 $2_1$ 2 | P 2 2 $2_1$ |
| a (Å) | 67.49 | 67.8 |
| b (Å) | 73.76 | 75.02 |
| c (Å) | 127.28 | 125.78 |
| Resolution range (Å) | 39.25 - 1.89 | 50.31 - 1.79 |
| Total reflections measured | 707,104 | 826,980 |
| Unique reflections | 100,181 | 61,413 |
| Completeness (%) (last shell) | 100 (100) | 99.8 (95.7) |
| I/$\sigma$ (last shell) | 10.4 (0.3) | 10.5 (0.4) |
| R(merge)(%) (last shell) | 9.3 (329.2) | 0.125 (4.902) |
| B (iso) from Wilson (Å2) | 35.3 | 27.68 |
| **Refinement statistics** | | |
| R-factor$^b$ (%) | 0.206 | 20 |
| R-factor$^c$ (%) | 0.247 | 23.4 |
| RMSD bond lengths (Å) | 0.0121 | 0.013 |
| RMSD bond angles (°) | 1.806 | 1.795 |
| **Ramachandran Plot statistics** | | |
| Favored region (%) | 93 | 96.68 |
| Allowed region (%) | 6.4 | 2.65 |
| Disallowed region (%) | 0.6 | 0.66 |

**Table S2. Residues comprising the dGFP-diS2 dimer interface as determined using PISA [2]**

<table>
<tr><th colspan="3">Chain A</th><th colspan="3">Chain B</th></tr>
<tr><th>Residue</th><th>H-bond</th><th>BSA (Å²)[a]</th><th>Residue</th><th>H-bonds</th><th>BSA (Å²)[a]</th></tr>
<tr><td>T38</td><td></td><td>0.99 |</td><td>T38</td><td></td><td>1.11 |</td></tr>
<tr><td>N39</td><td>H</td><td>33.64 |||||</td><td>N39</td><td>H</td><td>33.11 |||||</td></tr>
<tr><td>R73</td><td>H</td><td>90.31 |||||||||</td><td>R73</td><td>H</td><td>90.49 |||||||||</td></tr>
<tr><td>P75</td><td></td><td>35.66 |||||||||</td><td>P75</td><td></td><td>35.66 |||||||||</td></tr>
<tr><td>H77</td><td></td><td>23.80 |||||</td><td>H77</td><td></td><td>24.25 |||||</td></tr>
<tr><td>M78</td><td></td><td>0.83 |||||||||||</td><td>M78</td><td></td><td>0.83 ||||||||||</td></tr>
<tr><td>N146</td><td>H</td><td>9.04 ||</td><td>N146</td><td>H</td><td>9.17 ||</td></tr>
<tr><td>S147</td><td></td><td>3.48 ||</td><td>S147</td><td></td><td>3.36 ||</td></tr>
<tr><td>C148</td><td></td><td>55.15 |||||</td><td>C148</td><td></td><td>55.48 |||||</td></tr>
<tr><td>N149</td><td></td><td>20.93 |||||</td><td>N149</td><td></td><td>20.54 |||||</td></tr>
<tr><td>Y200</td><td>H</td><td>27.14 ||||||</td><td>Y200</td><td>H</td><td>27.09 ||||||</td></tr>
<tr><td>S202</td><td>H</td><td>24.80 ||||||||</td><td>S202</td><td>H</td><td>24.93 ||||||||</td></tr>
<tr><td>Q204</td><td>H</td><td>90.68 |||||||||</td><td>Q204</td><td>H</td><td>90.41 |||||||||</td></tr>
<tr><td>S205</td><td></td><td>5.11 |||</td><td>S205</td><td></td><td>5.20 |||</td></tr>
<tr><td>F206</td><td></td><td>107.03 ||||||||||</td><td>F206</td><td></td><td>107.32 ||||||||||</td></tr>
<tr><td>L207</td><td></td><td>16.74 ||||||||</td><td>L207</td><td></td><td>17.00 ||||||||</td></tr>
<tr><td>S208</td><td></td><td>37.49 |||||||</td><td>S208</td><td></td><td>37.31 |||||||</td></tr>
<tr><td>K209</td><td></td><td>3.69 |</td><td>K209</td><td></td><td>3.81 |</td></tr>
<tr><td>V219</td><td></td><td>7.83 ||</td><td>V219</td><td></td><td>8.00 ||</td></tr>
<tr><td>L221</td><td></td><td>34.80 |||||||</td><td>L221</td><td></td><td>34.97 |||||||</td></tr>
<tr><td>F223</td><td></td><td>68.67 |||||||||</td><td>F223</td><td></td><td>67.88 |||||||||</td></tr>
<tr><td>V224</td><td></td><td>0.68 |</td><td>V224</td><td></td><td>0.68 |</td></tr>
<tr><td>T225</td><td></td><td>29.93 ||||||||</td><td>T225</td><td></td><td>30.09 ||||||||</td></tr>
<tr><td>A226</td><td></td><td>0.48 |||||||||||</td><td>A226</td><td></td><td>0.36 ||||||||</td></tr>
<tr><td>A227</td><td></td><td>33.71 ||||||||||</td><td>A227</td><td></td><td>33.88 ||||||||||</td></tr>
<tr><td>G228</td><td></td><td>4.91 |</td><td>G228</td><td></td><td>4.46 |</td></tr>
<tr><td>I229</td><td></td><td>3.54 ||</td><td>I229</td><td></td><td>3.78 ||</td></tr>
<tr><td>T230</td><td></td><td>49.18 |||||</td><td>T230</td><td></td><td>48.90 |||||</td></tr>
<tr><td>M233</td><td></td><td>90.80 ||||||</td><td>M233</td><td></td><td>90.46 ||||||</td></tr>
</table>

[a], Buried surface area. Each vertical line represents percentage of buried surface area, with one bar equal to 10%.

**Table S3. H-bonds at the dGFP-diS2 dimer interface as determined by PISA[2]**

| Chain A | | | Chain B | |
|---|---|---|---|---|
| **Residue** | **Atoms[a]** | **Distance (Å)** | **Atoms[a]** | **Residue** |
| N39 | O | 2.95 | NH1 | N73 |
| R73 | NH1 | 2.95 | O | N39 |
| N146 | OD1 | 3.89 | OH | Y200 |
| Y200 | OH | 3.89 | OD1 | N146 |
| Y200 | OH | 2.16 | OE1 | N204 |
| S202 | OG | 3.37 | OG | N202 |
| Q204 | OE1 | 2.16 | OH | Y200 |

a, atoms are labelled by their PDB nomenclature.

**Table S4. Change in emission ratio of GFPdiS2-ZIP in the presence of various metal ions**

| Metal | Ex 483 nm:Ex 388 nm | | $R/R_0$ |
|---|---|---|---|
| | Before[a] | After[b] | |
| $CaCl_2$ | 0.61 | 0.71 | 1.16 |
| $MgCl_2$ | 0.65 | 0.69 | 1.05 |
| $MnCl_2$ | 0.75 | 0.85 | 1.13 |
| $ZnCl_2$ | 1.15 | 1.18 | 1.03 |
| $ZnSO_4$ | 1.12 | 1.12 | 1.00 |
| $Fe^{3+}$: $FeCl_3$[b] | 2.04 | 2.49 | 1.22 |
| $Fe^{2+}$: $FeSO_4$[b] | 4.26 | 4.08 | 0.96 |
| $CoCl_2$ | 0.66 | 0.77 | 1.16 |

a, the emission ratio immediately after (<30 sec) of 1mM metal ion.
b, 1 hr after addition.
c, higher ratio due to increase in emission on excitation at the higher wavelength due to the metal ion itself.

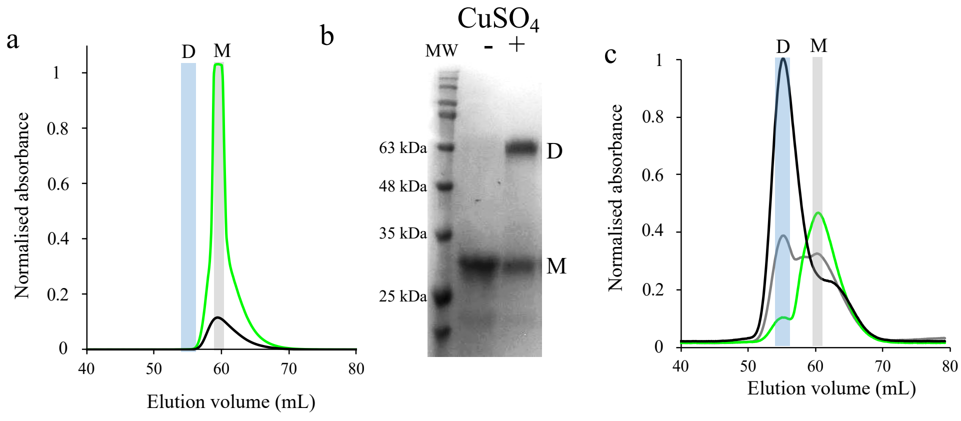


**Figure S1.** Dimerisation characteristics of GFP-diS1. (a) Size exclusion chromatography (SEC) of purified GFP-diS1. (b) Non-reducing SDS-PAGE of GFPdiS1 in the absence (-) and presence (+) 1 mM $CuSO_4$. The lower band present just below 25 kDa is commonly observed in fluorescent proteins due to cleavage in the chromophore on heat denaturation [3,4]. (c) Separation of dimer and monomer species by SEC after incubation with $CuSO_4$. On all figures, D signifies dimer and M monomer. Green, black and grey lines in the SEC elution profiles represent absorbance at 400 nm, 485nm and 280 nm, respectively.

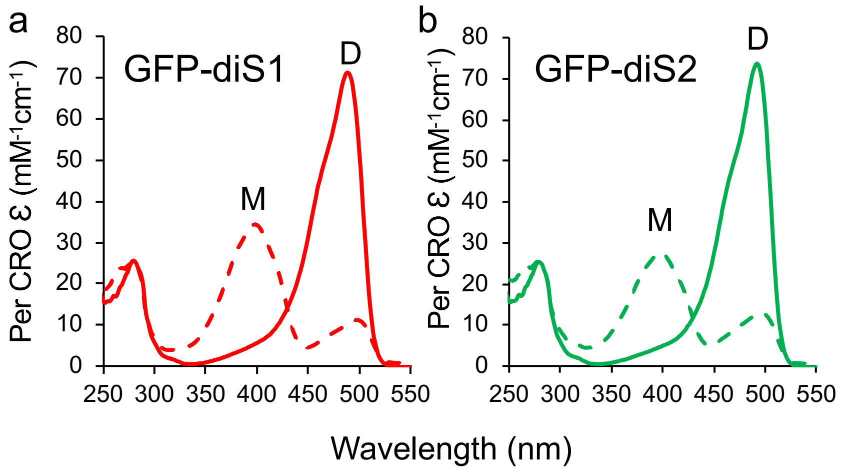


**Figure S2.** Absorbance spectra of (a) GFP-diS1 and (b) GFP-diS2 in their monomer (dashed lines) or dimer (solid lines) forms. Absorbance is shown as the per chromophore molar absorbance coefficient for direct comparison. . SEC was performed with either HiLoad™ 16/600 Superdex™ S75 pg.

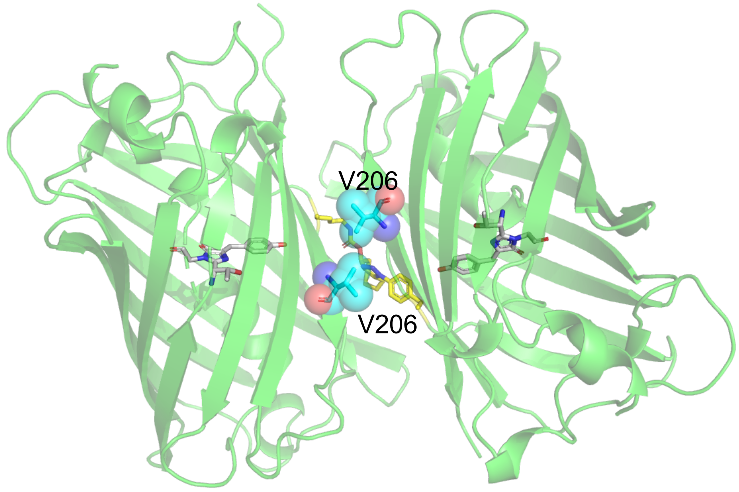


**Figure S3.** The position of V206 at the dimer interface of a sfGFP homodimer linked by alkyne-azide crosslink at residue 148 (PDB: 5nhn) [5].

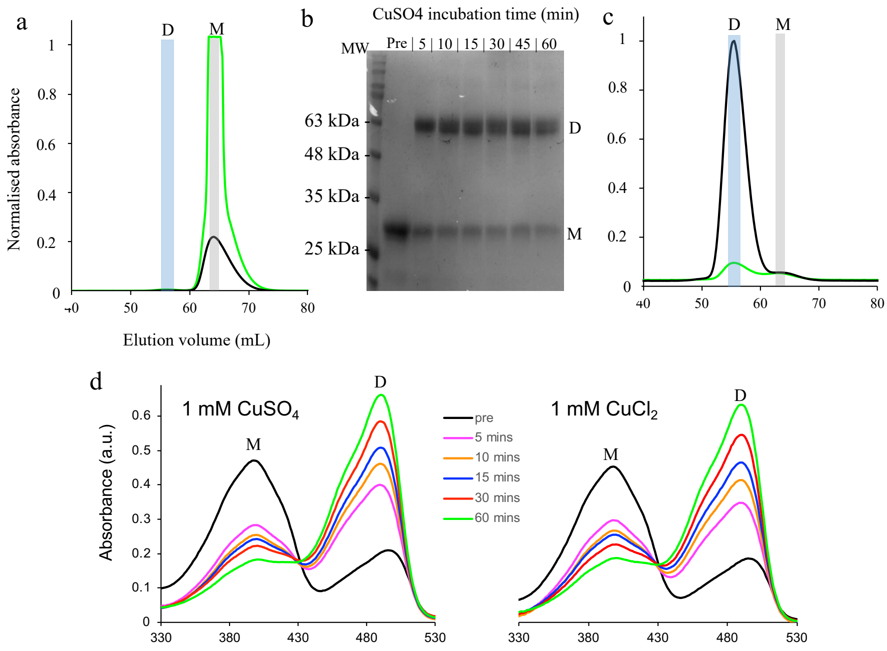


**Figure S4.** Dimerisation characteristics of GFP-diS2. (a) Size exclusion chromatography (SEC) of purified GFP-diS2. Green and black lines in the SEC elution profiles represent absorbance at 400 nm and 485nm, respectively. (b) Non-reducing SDS-PAGE of GFP-diS2 on addition of 1 mM $CuSO_4$. Samples were taken periodically after $CuSO_4$ addition. "Pre" signifies prior to $CuSO_4$ addition. The lane numbers are time of incubation in minutes. (c) Separation of dimer and monomer species by SEC after incubation with $CuSO_4$. (d) Change in absorbance spectra on addition of either 1 mM $CuSO_4$ or 1 mM $CuCl_2$ to 10 μM GFP-diS2. On all figures, D signifies dimer and M monomer. SEC was performed with HiLoad$^{TM}$ 16/600 Superdex$^{TM}$ S200 pg (Cytiva).

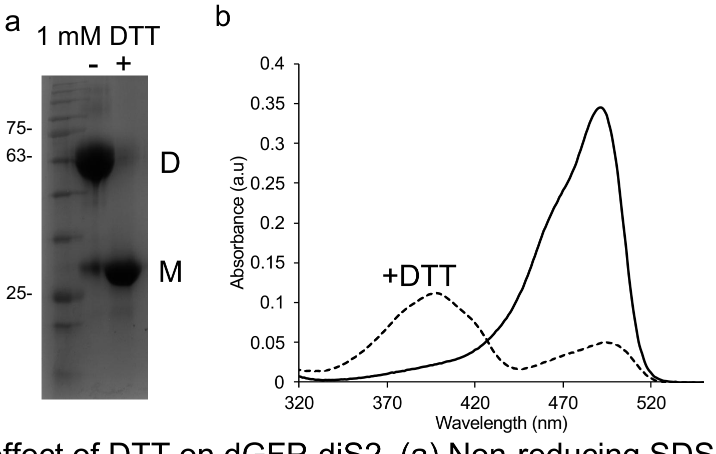


**Figure S5**. The effect of DTT on dGFP-diS2. (a) Non-reducing SDS-PAGE of the dimer before (-) and after (+) incubation of dGFP-diS2 for 5 min in 1 mM DTT. (b) Corresponding absorbance spectra (5 μM protein) before (solid line) and after (dashed line) incubation for 5 min with 1 mM DTT.

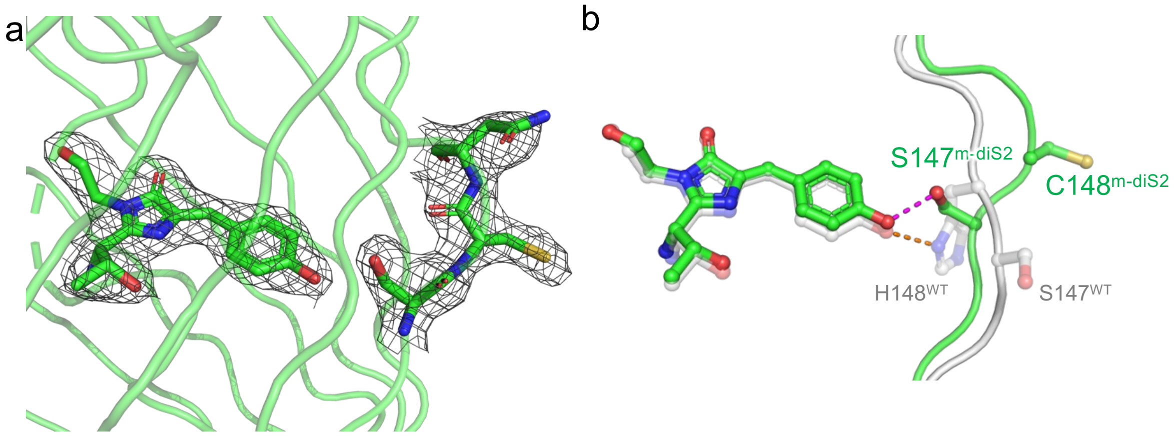


**Figure S6.** (a) The $2F_0$-$F_c$ map ($\sigma$ 1.0) of the chromophore and residues S147 and C148 in mGFP-diS2. (b) Comparison of the chromophore and residues 147 and 148 in WT sfGFP (grey; PDB 2b3p [6]) and mGFP-diS2 (green).

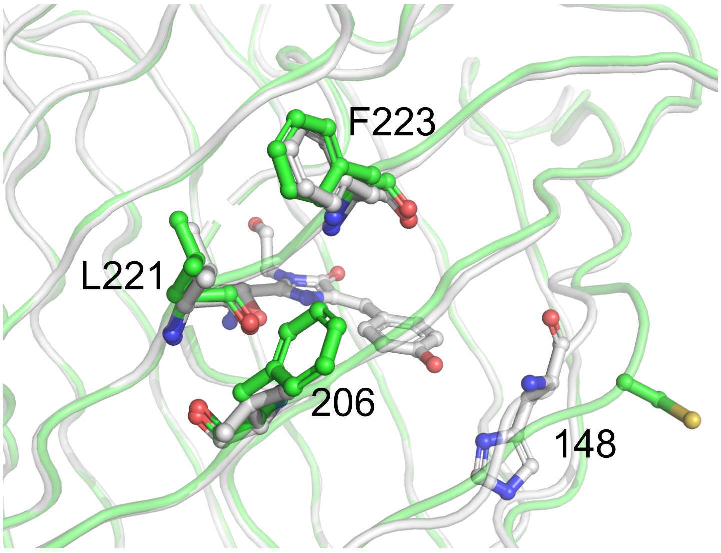


**Figure S7.** Structural comparison of WT sfGFP (grey) and mGFP-diS2 (green) centred around the V206F mutation.

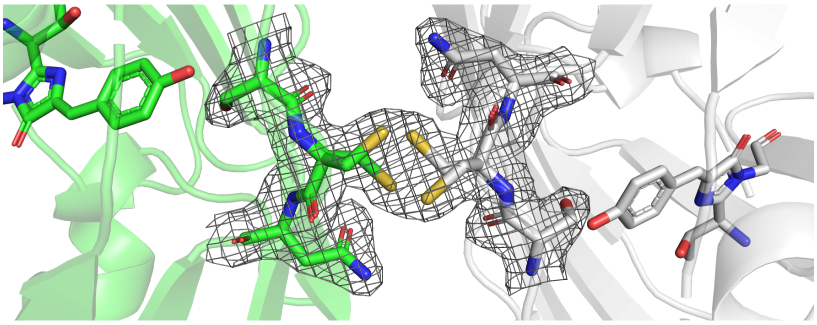

**Figure S8.** The $2F_0$-$F_c$ map ($\sigma$ 1.0) of the disulphide bridge region connecting each subunit (green and grey) of dGFP-diS2.

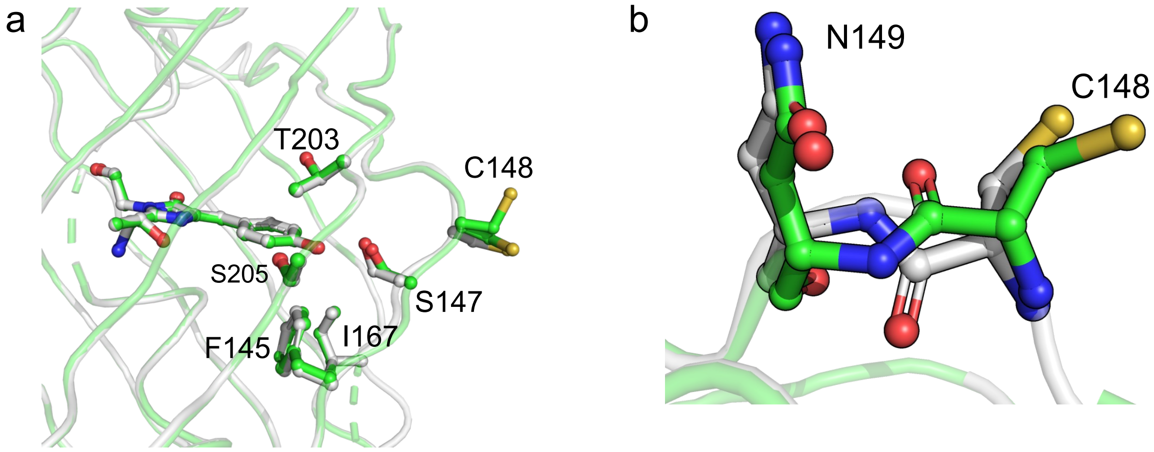


**Figure S9.** Structural overlay of mGFP-diS2 (grey) and dGFP-diS2 (green) showing (a) key residues in close proximity to the chromophore and (b) C148-N149 peptide bond plane flipping.

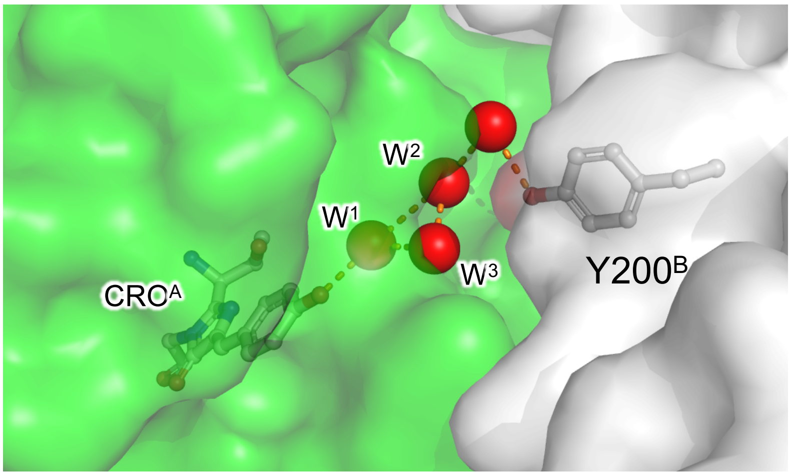


**Figure S10.** Surface view of dGFP-diS2 showing buried waters at the water dimer interface. Subunits A and B are coloured green and white respectively. Waters are shown as red spheres and labelled as in Figure 4 in the main text.

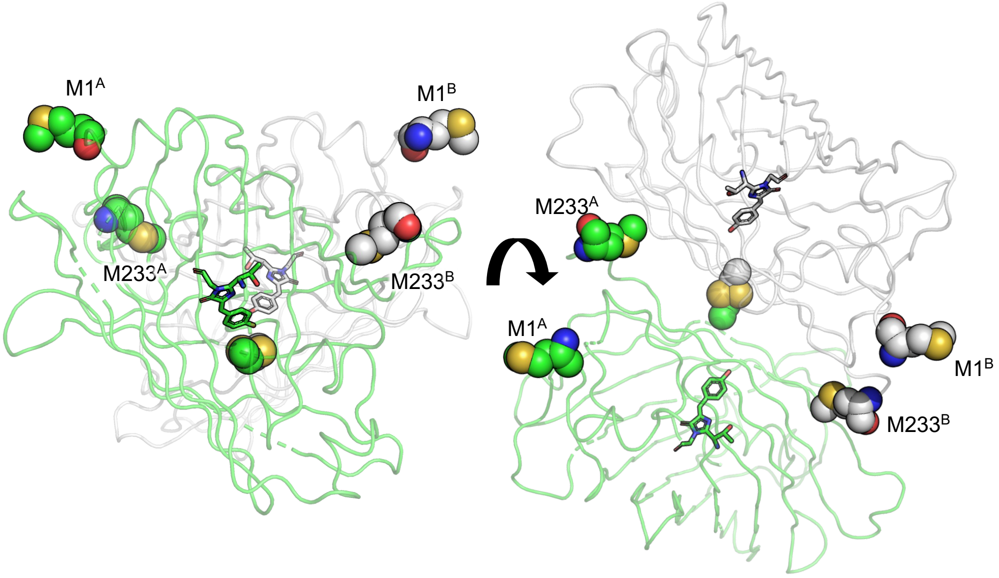


**Figure S11**. Location of N-termini (M1$^{A}$ and M1$^{B}$) and last observed C-terminal residue (M233$^{A}$ and M233$^{B}$) in each monomer unit in dGFP-diS2.

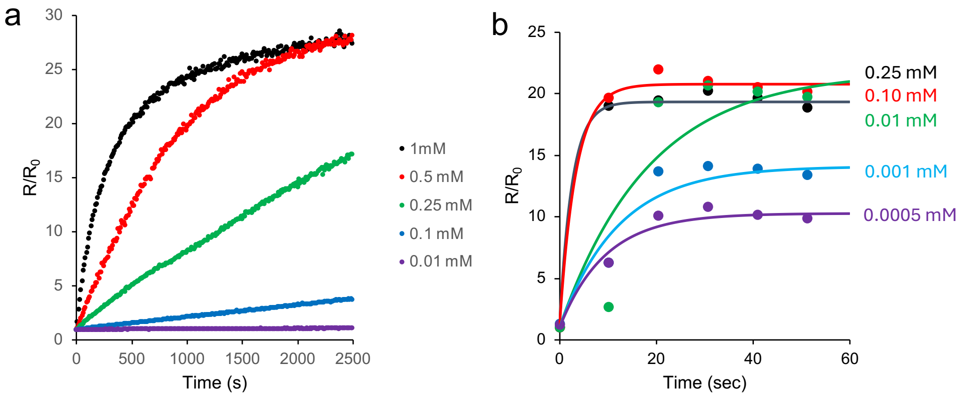


**Figure S12.** Rate of change in fluorescence emission ratio (Ex$^{CRO\text{-}O^-}$:Ex$^{CRO\text{-}OH}$) in the presence of difference concentrations of (a) $Cu^{2+}$ ($CuSO_4$) and (b) $Cu^+$ ($CuSO_4$ plus ascorbate). The copper concentration for each curve is shown on the plots themselves.

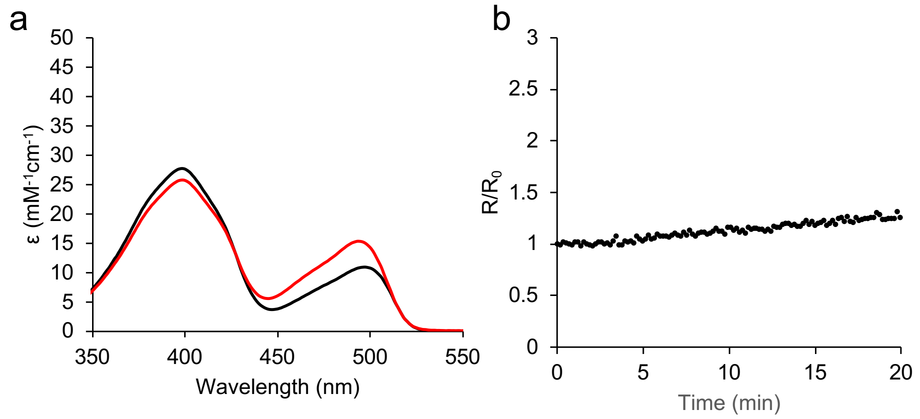


**Figure S13.** The effect of ascorbate on the spectral properties of GFP-diS2-ZIP. (a) The absorbance spectra of GFP-diS2-ZIP (5 μM) before the addition of 500 μM ascorbate (black) and after 1 hr incubation (red). (b) The fluorescence emission ratio ($Ex^{CRO-O-}$:$Ex^{CRO-OH}$) time course of 5 μM GFP-diS2-ZIP in the presence of 500 μM ascorbate.

**Supporting References.**


1 S. C. Reddington, E. M. Tippmann and D. D. Jones, *Chemical Communications*, 2012, **48**, 8419–8421.

2 E. Krissinel and K. Henrick, *J Mol Biol*, 2007, **372**, 774–97.

3 J. Wei, J. S. Gibbs, H. D. Hickman, S. S. Cush, J. R. Bennink and J. W. Yewdell, *Journal of Biological Chemistry*, 2015, **290**, 16431–16439.

4 H. S. Auhim, B. L. Grigorenko, T. K. Harris, O. E. Aksakal, I. V. Polyakov, C. Berry, G. dos P. Gomes, I. V. Alabugin, P. J. Rizkallah, A. V. Nemukhin and D. D. Jones, *Chem Sci*, 2021, **12**, 7735–7745.

5 H. L. Worthy, H. S. Auhim, W. D. Jamieson, J. R. Pope, A. Wall, R. Batchelor, R. L. Johnson, D. W. Watkins, P. Rizkallah, O. K. Castell and D. D. Jones, *Commun Chem*, 2019, **2**, 83.

6 J. D. Pédelacq, S. Cabantous, T. Tran, T. C. Terwilliger and G. S. Waldo, *Nat Biotechnol*, 2006, **24**, 79–88.
